# Nanoscale prognosis of colorectal cancer metastasis from AFM image processing of histological sections

**DOI:** 10.1101/2022.05.06.490873

**Authors:** Vassilios Gavriil, Angelo Ferraro, Alkiviadis-Constantinos Cefalas, Zoe Kollia, Francesco Pepe, Umberto Malapelle, Caterina De Luca, Giancarlo Troncone, Evangelia Sarantopoulou

## Abstract

Early ascertainment of metastatic tumour phases is crucial to improve cancer survival, formulate an accurate prognostic report of disease advancement and, most important, quantify the metastatic progression and malignancy state of primary cancer cells with a universal numerical indexing system. This work proposes an early improvement of cancer detection with 97 *nm* spatial resolution by indexing the metastatic cancer phases from the analysis of atomic force microscopy images of human colorectal cancer histological sections. The procedure applies variograms of residuals of Gaussian filtering and theta statistics of colorectal cancer tissue image settings. The methodology elucidates the early metastatic progression at the nanoscale level by setting metastatic indexes and critical thresholds from relatively large histological sections and categorising the malignancy state of a few suspicious cells not identified with optical image analysis. In addition, we sought to detect early tiny morphological differentiations indicating potential cell transition from epithelial cell phenotypes of low to high metastatic potential. The metastatic differentiation, also identified by higher moments of variograms, sets different hierarchical levels for the metastatic progression dynamic, potentially impacting therapeutic cancer protocols.

## Introduction

Tumour metastasis is the migration of cancer cells from the primary tumour cores to the lymph nodes, tissues, or distant organs. Metastasis is responsible for 90% of colorectal cancer (CRC) deaths; therefore, early diagnosis is critical for patient survival^1^. Metastasis is a complex process that involves morphological adjustments and the attachment of cancer cells to other cells and the extracellular matrix (ECM). It represents a key hallmark^2^ of malignance’s progression towards a higher pathological state. Therefore, indexing assessment of the metastatic state and its early prediction is fundamental for enlightening cancer progression, improving early cancer prognosis and developing therapeutic schemes^3,4^. Tissue microenvironmental factors, including stiffness and topography (nuclei’s shape, morphology, and texture specificity), contribute to the targeting preferences of metastatic cancers^5–9^ because biological and mechanical/topographic parameters are associated with cancer cell proliferation, migration and metastasis^7,10,11^. Cancer cells regulate their stiffness to match the ECM local environment by adjusting viability to different structural proteins’ complex ECM topographical environment.

Metastasis of variable percentage may arise in all stages^12^, indicating that common histological and cytological findings are necessary but insufficient to identify high-risk characteristics and predict metastatic phases. To improve patients’ survival, it is mandatory to ascertain the tumour stage accurately, formulate a universal prognostic report about disease progression, and, most importantly, identify the metastatic phase and heterogeneity of primary cancer cells as early as possible^13,14^. Even though the latest CRC TNM classification protocols of malignant in regional lymph nodes are considered, and each pathological stage is further subdivided^15^, early state and novel classification schemes are needed, and research aims to establish biomarkers for early and reliable tumour diagnosis and metastasis prognosis. The correlation between metastasis and tumour histological alterations was recognized in the early mid-nineteenth century. Since then, optical and electronic microscopy has been applied for routine cancer diagnostics by visual interpretation of ultra-thin, two-dimensional tissue sections, where histopathologists decide whether tissue regions are cancerous and classify the malignancy level^16^.

Still, diagnosis and classification of cancer are operator-dependent and thus imperilled to errors. Also, negative factors include the inherent limitations of magnification, the field of view, contrast, and small focal depth of optical systems^17^. Consequently, besides optical imaging state of the art histological image analysis software and texture algorithms, exploiting the microscopic variations of cells’ shapes and tissue morphologies are needed for early and reliable prediction of metastasis. Along the above lines, a novel methodology of probing the mechanics of tumours emerged as a supportive method to find the link between the mechanical properties of single tumour cells and their metastatic potential^18–20^. However, although several techniques exist, including the atomic force microscopy (AFM)^5,21,22^, to measure the mechanical properties of single cells, information on the mechanics of tumour cells in the ECM is missing because most measurements are made on cultured tumour cells^10,23^. Besides, each method has a particular set of parameters that do not consider patient-to-patient variations, which is an additional drawback in comparing different studies^5,24^.

Machine vision and learning methods were also applied as complementary approaches to microscopic histopathological examination and molecular-based approaches for cancer prediction and prognosis^25–27^. The established digital histopathology image analysis is based on tissue image classification and tiny segmented structures, including nuclei and cells^28^. However, machine learning still experiences numerous technical and organizational challenges and limitations because of the complexity of tissue morphology, tumour heterogeneity, and diversity of shape, location, and size of the tumour segmentation. Developing accurate and efficient algorithms is still a challenging issue^26^.

Likewise, mathematical modelling of complex natural systems, including tumours, aim to characterize architecture and decode spatial and temporal complexity and heterogeneity, commonly appearing in nature^29^. Fractality, complexity, and structure statistics discriminate tags suitably from Euclidean morphometric measurements (e.g. length, volume, density)^30–32^ and several methods^33^ developed to study physical entities in many different contexts^34–40^. For example, the generalised method of moments (GMM) is viewed as an extension of the *z*-height correlation functions. Variograms also are used extensively in geology and medicine^41–43^ to quantify images’ spatial variability and correlation distances. A variogram expresses the expected square difference between two data values separated by a distance-vector, e.g., grayscale values between pixels in optical microscopy or *z*-height values in AFM images. Overall, one or two-dimensional variograms (*1D, 2D*) are visual expressions of the spatial correlation of image points.

Variograms were used in diagnosis, including spatial tissue displacement of ultrasound elastography in areas surrounding needles^44^, image-guided neurosurgery^45^, and non-subjective evaluation of chromatin in cell proliferation and apoptosis^46^. Similarly, in magnetic resonance of *3D* brain structural changes^47^ and spatial autocorrelation stiffness differences between aortic and pulmonary valve interstitial cell^48^. Internal tension and sub-cellular spatial distribution differentiate metastatic and non-metastatic cells and tissues. Variograms applied to *2D* malignant breast tissue images^49^, anticancer treatment^50^ or differentiation between melanomas and normal skin tissues^51^.

Although low spatial resolution optical imaging^44^ utilizes variograms^42^, an early cancer prognostic tool implies tissue structural differentiation at the nanoscale level^52^. However, a reliable, label-free, non-invasive approach for identifying and quantifying nanoscale metastatic differentiation on conventional histological sections is challenging^53^. In this direction, AFM is suitable for non-destructive *3D* imaging of cells and tissues with nanometric resolution^19,54,55^. So far, few AFM studies have analyzed formalin-fixed and paraffin-embedded (FFPE) cancer histological tissues because of diagnostic and prognostic constraints^56,57^. Second-order effects and lacunarity (distribution, size of gaps between cells)^58^ were proposed as marking factors in histopathology image analysis. The correlation between the fractal dimension of AFM images and the *z*-scale factor served as a mechanical mark of human lung carcinoma^59^. Analysis of AFM adhesion of cells^60^ reveals that fractality differences are evident when premalignant cells transform into cancerous^61,62^.

Variogram analysis is based on the hypothesis that images’ statistical mean and variance are independent of pixels’ location. Also, statistical mean and variance commonly bear comparable values for entities belonging to the same hierarchical group, such as the different sets of metastatic and non-metastatic CRC AFM images of histological tissues. For example, the variograms of domain size Gaussian filtering (DSGF) differentiate similar but different hierarchies^43^.

In this work, small biological features discriminate AFM images of metastatic/non-metastatic CRC tissues. A significant sensitivity improvement in differentiating metastatic/ non-metastatic stages in CRC cells was obtained by applying moment variograms of residuals of Gaussian filtering and theta statistics^63^ in 50 *μm x* 50 *μm* AFM cancer histological images from three different patients. Likewise, AFM image theta statistics incorporate inclination histograms of tiny planar segments of CRC histological sections. The theta distribution skewness could differentiate the signatures of different hierarchical groups as metastatic and non-metastatic tissues.

Furthermore, towards establishing early quantifying markers of metastatic phases, the differentiation between metastatic and non-metastatic tissues was approached with rescaled range, surface statistics, and phase analysis in AFM imaging. Results were compared with those from variograms and theta statistics. Noticeably, the novelty and state-of-the-art of the current work are grounded on improving metastatic differentiation by higher moments of variograms.

The tactic aims to provide an insight into the metastatic hierarchical levels and the dynamics of metastatic evolution by early diagnosing the malignancy condition of suspicious cells (typically a few) not identified by optical microscopy when subtle signs appear. We sought to identify early tiny morphological changes indicating potential cell transition from an epithelial phenotype typical of cells with a low metastatic potential to a mesenchymal phenotype that marks high mobility cell features and provides quantifying universal metastatic indexes and critical thresholds.

## Results

### Optical and AFM Microscopy of CRC Histological Sections

Typical AFM CRC metastatic and non-metastatic histological tissue images, extracted during 2021 from three patients, are shown in Fig. 1.

**Fig. 1.**
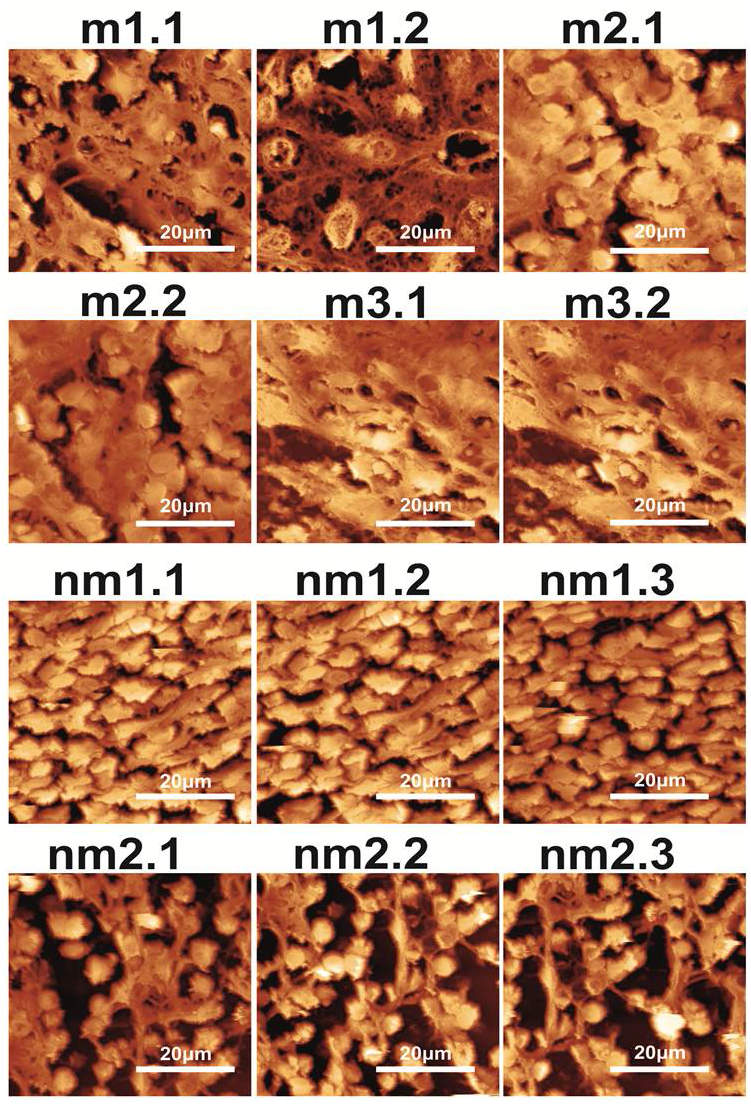
AFM images of metastatic (m1.1-m3.2) and non-metastatic (nm1.1-nm2.3) human CRC histological sections. The first and second numbers refer to the patient and sample, respectively.

The first and second indexing numbers are associated with the patient and sample parts. In AFM images, the visual differentiation between the two classes of metastatic and non-metastatic tissues is unclear. On the contrary, optical images (*4x, 20x*, 40*x*) of hematoxylin stained CRC histological sections unveil metastatic/non-metastatic differentiation, Fig. 2. The cells of metastatic tissues, Figs. 2a,2b, are closely spaced compared with the non-metastatic ones, Figs. 2d,2e. Nevertheless, optical microscope differentiation between metastatic and non-metastatic cells might be subjective and, in some cases, depends on the operator.

**Fig. 2.**
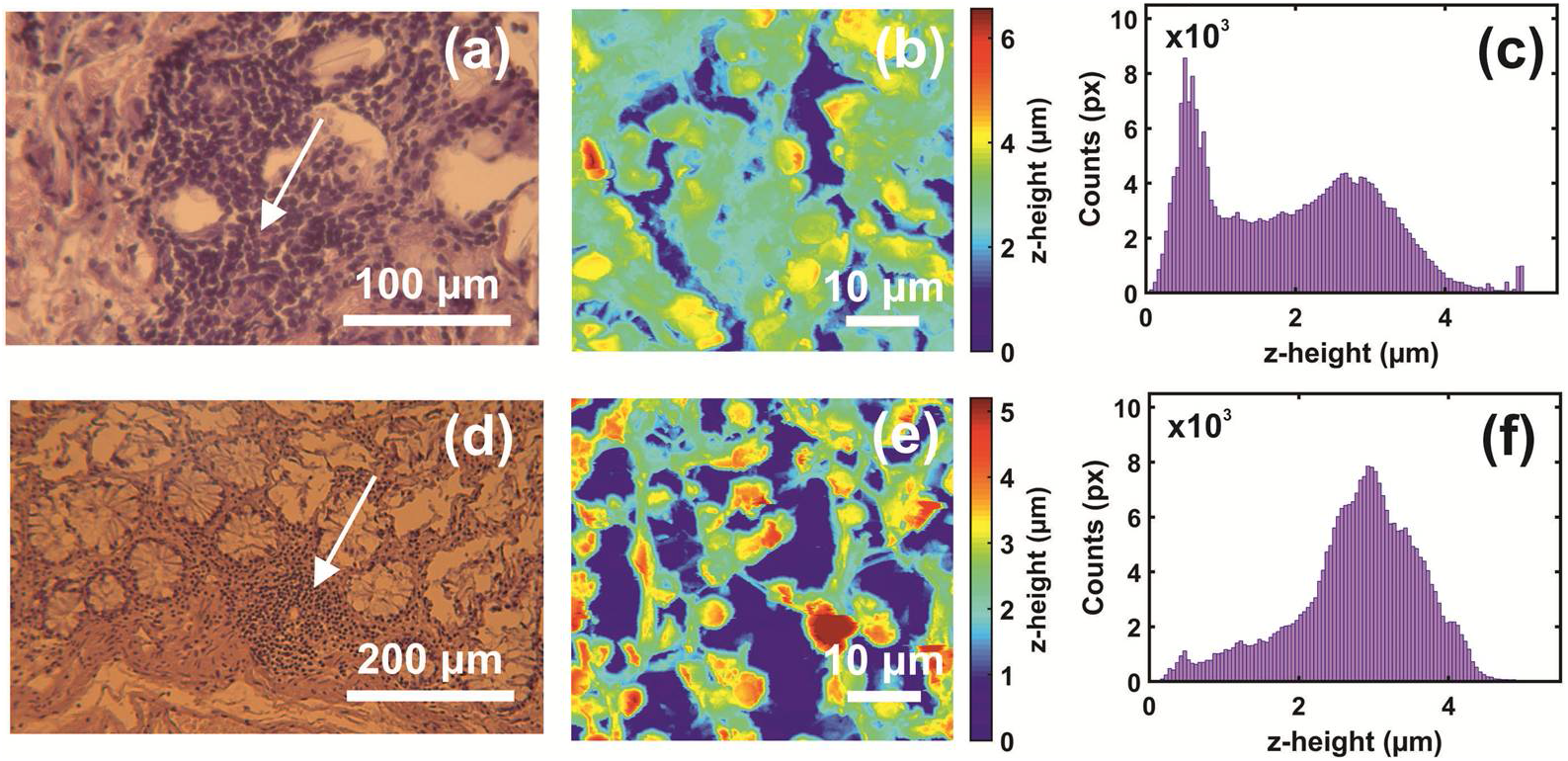
Optical, AFM images and *z*-height distribution of human CRC histological metastatic/non-metastatic sections. **(a)** Optical image (40 *x*) of a metastatic section (0.17 *mm x* 1.73 *mm*). **(b)** AFM image of metastatic tissue at the point a of the image (a) (arrow). **(c)** The *z*-height distribution from the AFM image of image (b). **(d)** Optical image (20*x*) of non-metastatic histological section. **(e)** Non-metastatic AFM of image (d) (arrow). **(f)** The *z*-height distribution from the AFM image (e).

### Variograms of Residuals Gaussian Filtering

The two dimensional (*2D*) variograms of the residuals of the Gaussian filtered AFM images, Fig. 1, of metastatic and non-metastatic histological tissues along with all directions sustain closed elliptic and open contours, Fig. 3. The same colour closed areas characterize equal RMS deviations of small size spatial scale differences (small lag vectors). Spatial correlations with dimensions below 0.5 *μm* are from small biological and structural tissue topologies. For a Gaussian filter applied with standard deviation *σ (px)*, the kernel box size along each axis is 6*σ*+1*(px)*, and the lag vectors’ zero limits (nugget) is 1 *px*, Fig. 4a. For standard deviation *σ* values between 2.5, 5, and 10 *px*, the magnitude of RMS deviation of closed contour areas diverges for metastatic and non-metastatic phases, Fig. 3, Supplementary Fig. S1 online. Close to the centre of the *2D* variograms, a relatively large RMS deviation is the typical signature of non-metastatic tissues. The colour indexing reveals that the mean RMS deviation of the metastatic and non-metastatic tissues is ~0.17 and ~0.27 *μm* (blue-arctic, yellow-lemon, Fig. 3, respectively). Therefore, the RMS deviation of the non-metastatic phase is noticeably larger than the metastatic one. The differentiation between metastatic and non-metastatic tissues is also retained for lower resolutions images of equal size (50 *μm x* 50 *μm*), e.g. for 256 *px x* 256 *px*, and 128 *px x* 128 *px* image sizes and *σ* between 2.5 and 10 *px*, Supplementary Fig. S1 online.

**Fig. 3.**
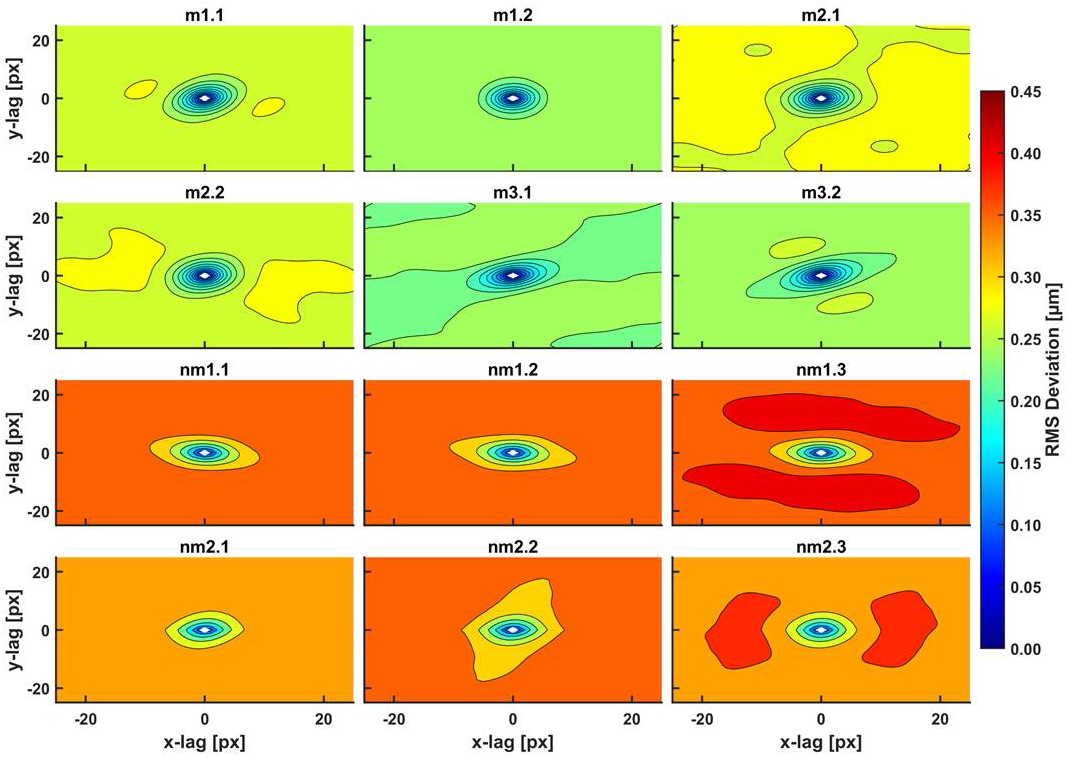
*2D* RMS deviation spectra of metastatic/non-metastatic CRC histological sections of the AFM images, Fig. 2. The spectra were taken with a *3D* Gaussian high pass filter. The RMS deviation images represent a statistical measure of the deviation of heights within an area at a particular scale. The plot shows elliptical-like contours for small scales (100 *nm*-1 *μm*, 1-10 *px*) of equal-value RMS deviation for a given colour. Different RMS deviations from the colour index are noticed for metastatic/non-metastatic sections. Non-metastatic CRC sections are characterized by higher values of RMS deviation within the closed and open areas compared to non-metastatic ones.

**Fig. 4.**
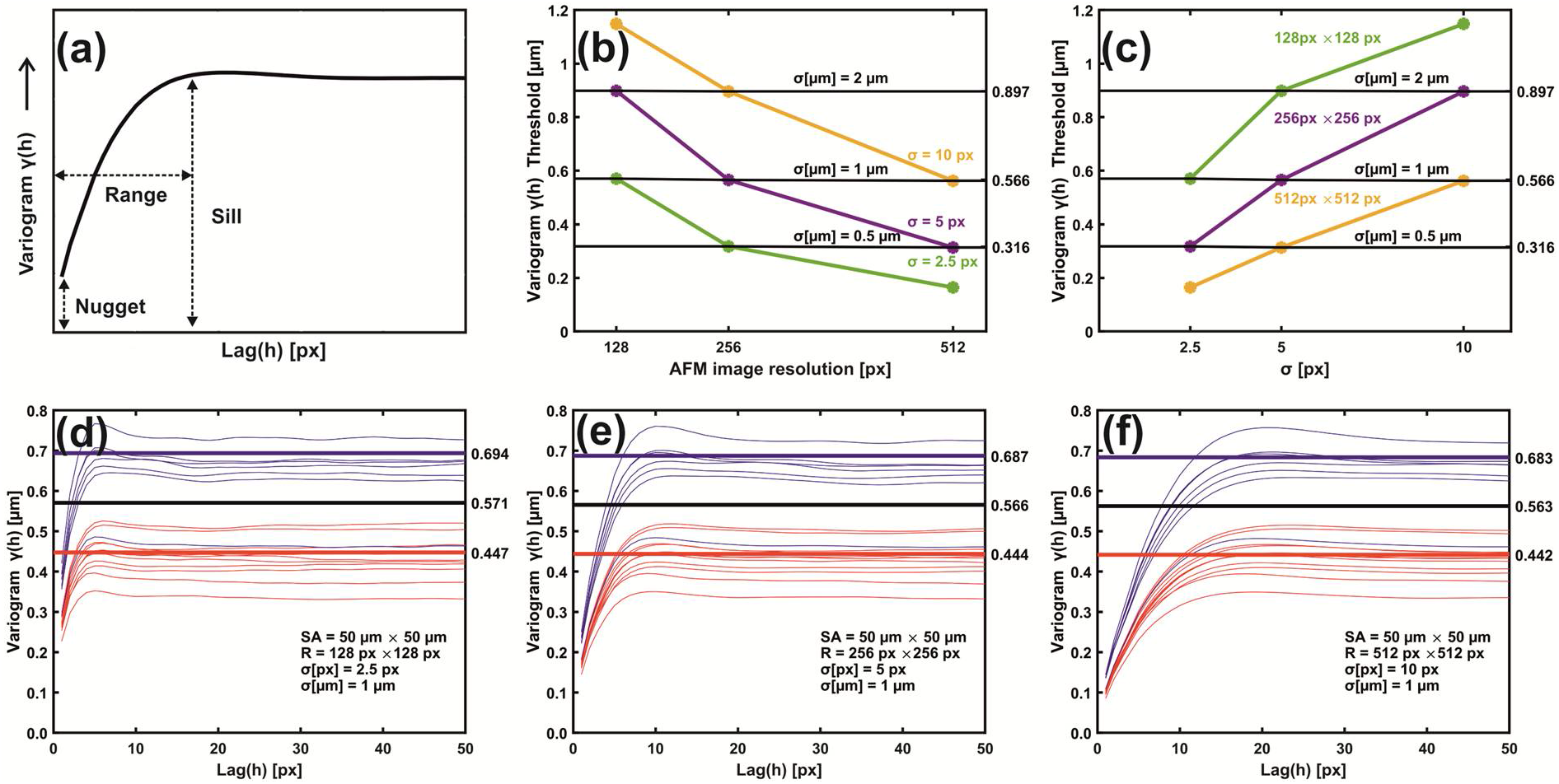
Metastatic/non-metastatic *1D* variograms of residuals Gaussian filtering. **(a)** Variogram’s typical parameters. **(b)** Metastatic threshold *vs*. AFM image resolution for different σ *[px,μm]*. The metastatic threshold stays invariant for identical σ *[μm]* values. **(c)** Metastatic threshold *vs*. σ[*px*] at different AFM image resolutions. The metastatic threshold is invariant for identical σ (*μm*) values. **(d)** Variogram lines of metastatic (red) and non-metastatic (blue) groups. AFM image size 50 *μm x* 50 *μm*, resolution 128 *px x* 128 *px*, standard deviation *σ [μm]*=1 *μm*, *σ [px]=2.5 px*. The black sill line is the metastatic threshold defined as the median value of mean sill values of metastatic and non-metastatic tissues. Above the threshold line tissues are non-metastatic and metastatic below the line. Red and blue lines are the mean metastatic/non metastatic sill values. **(e)** The same as (d) with image resolution 256 *px x* 256 *px* and *σ [px]*= 5 *px*. **(f)** The same as (e) with image resolution 512 *px x* 512 *px*, and *σ [px]*= 10 *px*.

Comprehensive interpretation and quantification of *2D* metastatic and non-metastatic variograms are gained by *1D* variograms, Figs. 4a–4f. The amplitude of lag vectors for all directions is along the *x*-axis. Along the *y*-axis, the non-overlapping sill values *γ(h)* of metastatic (red curves) and non-metastatic (blue curves) histological tissues represent lag vectors of zero correlation, with a relatively wide gap between the sill values of the two histological groups. Also, the sill values of non-metastatic tissues are constantly placed above the sill values of metastatic ones, Figs. 4d–4f. Most importantly, for different image resolutions (pixels per line) and the same σ (*μm*), the sill indexes are invariant for the metastatic and non-metastatic groups, Figs. 4d–4f, Supplementary Fig. S2 online. The mean sill value of each metastatic and non-metastatic group (red and blue lines parallel to the *y*-axis) is extracted from the average sill values of the associated histological tissues, Figs. 4d–4f. The median value of mean sill values of metastatic and non-metastatic tissues defines threshold lines (black line) above which tissues are non-metastatic and metastatic below the line. For different image resolutions and identical *σs*, equal to 1 *μm*, the set of three different pixel and *σ* pairs, (128 *px x* 128 *px*, 2.5 *px* (1 *μm*)), (256 *px x* 256 *px*, 5 *px* (1 *μm*)), (512 *px x* 512 *px*, 10 *px* (1 *μm*)), retains almost a constant threshold sill value, equal to 0.571, 0.566 and 0.563 *μm*, Figs. 4d–4f.

Similarly, two sets of pixel and identical *σ* (*μm*) values ((128 *px x* 128 *px*, 5 *px (2 μm)*, (256 *px x* 256 *px*, 10 *px* (*2 μm*)), and ((256 *px x* 256 *px*, 2.5 *px* (*0.5 μm*), (512 *px x* 512 *px*, 5 *px* (*0.5 μm*)) retain almost the same threshold sill values, equal to 0.899, 0.896, and 0.318, 0.314 *μm*; Supplementary Fig. S2 online. Variograms of low-resolution images and large *σ’s* bear wider gaps and high uncertainty between the mean sill values of metastatic and non-metastatic variogram groups (bands), Supplementary Fig. S2 online. Relatively large *σs* amplify the uncertainty of information. The optimum metastatic differentiation for the current experimental configuration is obtained for a resolution of 512 *px x* 512 *px* and *σ*=5 *px*. The threshold criteria for differentiating metastatic and non-metastatic tissues were successful in 17 out of 18 samples, except sample nm2.4, which is non-metastatic, but appeared to have metastatic behaviour. However, by applying higher moments than 2 (*vide infra*), the nm2.4 sample has the correct non-metastatic behaviour.

### Moments of Gaussian Filtering Residuals Variogram

Gaussian filtering residuals variograms of higher moments upsurge the differentiation between metastatic and non-metastatic AFM images. For large scaling exponents *q*, the difference between metastatic and non-metastatic tissues further widens than the lower *q* values, Figs. 5a–5d. For example, the variogram sill value (512 *px x* 512 *px*, *σ*=5 *px*) for *q>3* is always higher than *q<3* in all non-metastatic samples compared to the metastatic ones, Figs. 5a–5d, Supplementary Fig. S3 online. Furthermore, the nm2.4 tissue sample, the unsuccessful exception in the *1D* variograms threshold criterion that behaves as a metastatic one, now adopts the correct non-metastatic behaviour for higher moments (*q>2*), agreeing with the pathologist’s examination. However, for different image resolutions and Gaussian filtering *σ*, the moments that give the corrected result for the nm2.4 tissue deviate, Supplementary Fig. S4 online. Therefore, the threshold criterion of metastasis varies for different moments. Consequently, the metastatic differentiation is improving at higher moments.

**Fig. 5.**
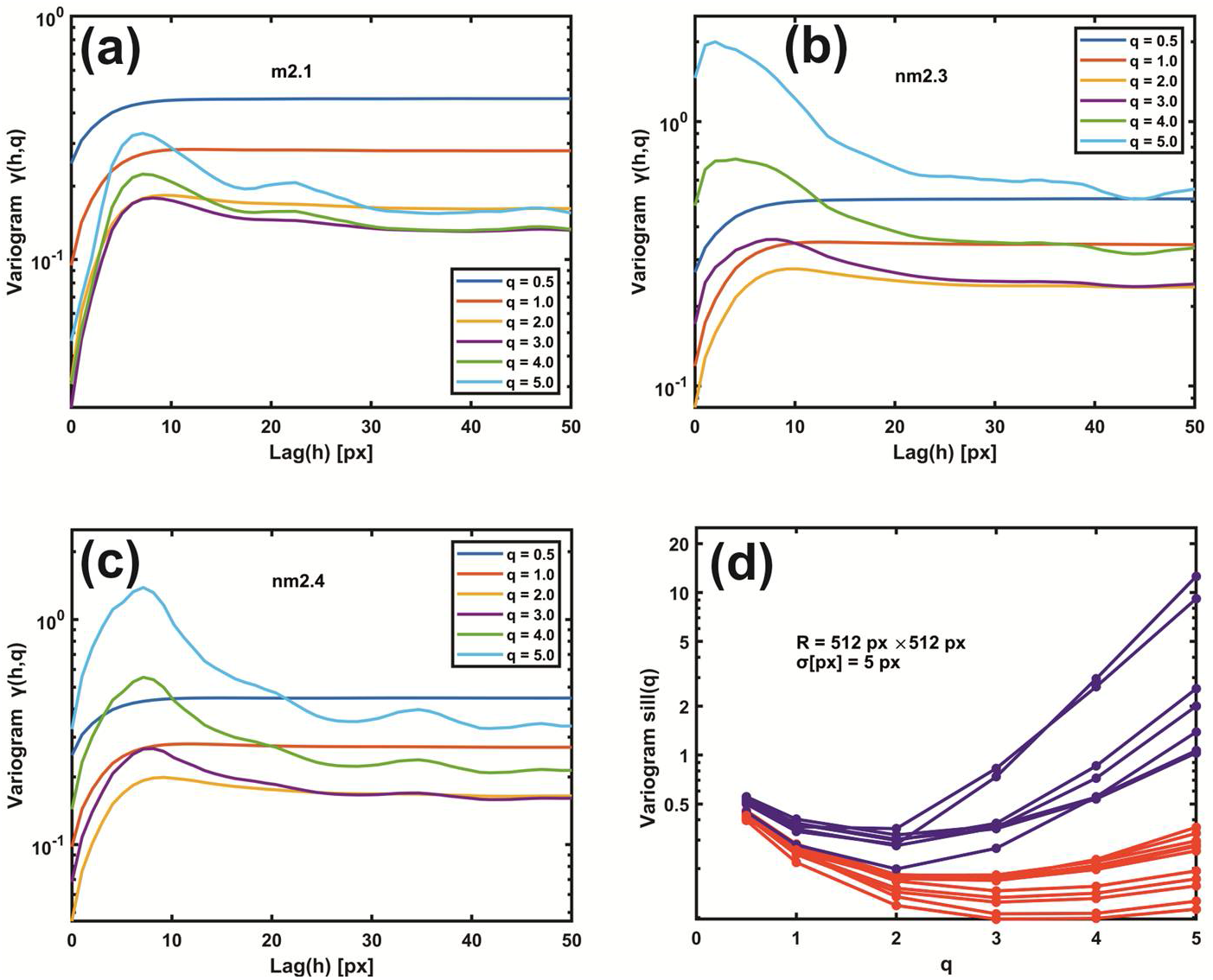
Gaussian filtering residuals variograms of different moments *q*. **(a)** Gaussian filtering residuals variogram of moments *q* from 0.5 to 5.0 for the metastatic tissue m2.1. **(b)** The same as (a) for the metastatic tissue nm2.3. **(c)** The same as (b) for the non-metastatic tissue nm2.4. Threshold criteria for differentiating metastatic and non-metastatic tissues, Fig. 4, are not functioning for the sample nm2.4. **(d)** Gaussian filtering residuals variograms of higher moments upsurge the differentiation between metastatic (red) and non-metastatic (blue) groups of lines. For higher moments than 2, the nm2.4 tissue sample has the correct non-metastatic state.

### Theta Statistics

Differences in the theta distribution profiling may be critically associated with different biological interactions between metastatic tumour cells and the ECM, leading to tissue differentiation. Other surface roughness characteristics in metastatic tissues (11 tissue samples) lead to notably broader inclination-angle distributions than the non-metastatic ones (7 tissue samples). The last is characterized by sharp peaks in the theta distribution diagram. It appears that non-metastatic tissues are typified by a structural surface regularity, highlighted by the sharp peaks at higher theta values, Fig. 6a. In contrast, random patterns and de-oriented structures define the metastatic phase. Skewness and kurtosis are differentiating measures in theta distribution. The skewness, Fig. 6b, of theta distribution of all metastatic sample AFM images is positive, agreeing with Fig. 6a.

**Fig. 6.**
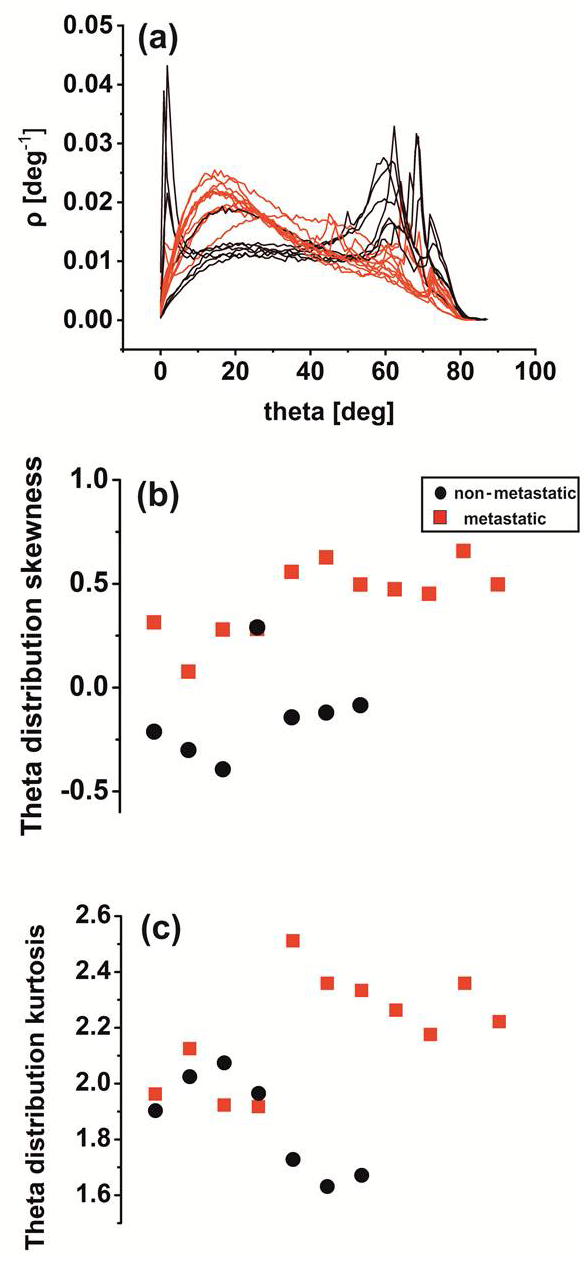
Theta statistics of metastatic/non-metastatic CRC histological sections. **(a)** Theta spectra of metastatic (red)/non metastatic (black) sections. The metastatic sections have one *maximum* value at 15°. The non-metastatic sections have two *maxima*, the first at small angles (2°) and the second at high angles (>60^0^). **(b)** Theta distribution skewness of metastatic/non-metastatic CRC histological sections with negative and positive skewness values. **(c)** Theta distribution kurtosis of metastatic/non-metastatic CRC histological sections.

In contrast, the skewness of non-metastatic tissues is negative (except for one sample), owing to the sharp peaks on the right side of the graph. In addition, kurtosis of theta distribution deviates from zero in all AFM images for metastatic and non-metastatic samples, leading to non-normal distributions, as expected, Fig. 6c. Although not in all cases, the skewness and kurtosis of metastatic tissues tend to have relatively large values.

### Surface Analysis

The standard statistical parameters of stained CRC histological sections of AFM images were calculated; Supplementary Figs. S5a,S5b online. The *z*-height distribution values of the AFM images of metastatic CRC histological tissues appear to have a wider dispersion around a mean value and obtain far extremer values than the non-metastatic ones; Supplementary Fig. S5a online. Also, the RMS roughness values, red squares, of metastatic tissues are smaller than circular black values for non-metastatic ones; Supplementary Fig. S5b online. Contrary to *1D*, *2D* variograms, and theta distribution, the surface analysis does not clearly distinguish between metastatic and non-metastatic phases. However, there is an underlying tendency of lower roughness values for metastatic tissues compared to non-metastatic ones, in agreement with the results from variograms and theta analysis, Figs. 3,4.

### Rescaled Range Analysis (Hurst exponent)

Rescaled range analysis/surface statistics is also applied along the same direction for 512 lines of each tissue image; Supplementary Figs. S5c-S5e online. First, each of the *2D* AFM images was transformed into a *1D* array by putting every line of 512 *px* one after another, and the Hurst exponent^64^ of each 512 *px* x 512 *px* array string was calculated, Supplementary Fig. S5c online. The same analysis was also performed for every line of an AFM image. Then, the mean value of the Hurst exponent of each AFM image was calculated for all lines, and the histogram was plotted; Supplementary Fig. S5d online.

The Hurst exponents and their trends extracted from the two algorithms are dissimilar; Supplementary Figs. S5c,5d online. The differentiation is expected because the two methods bear different correlations and connectivity between lines. The distribution histogram of the Hurst exponent distributed between the 512 lines is shown in Supplementary Fig. S5e online. There is a considerable variation of the Hurst exponent with the number of lines. The differentiation between metastatic and non-metastatic tissues is unclear, despite shifting the distribution function to the right relative to the *maximum* mean value for the metastatic tissues. The Hurst exponent does not differentiate between metastatic and non-metastatic tissues. Rescaled range analysis as second-order statistics usually provides insights for monofractal systems. However, metastasis is a dynamic process that drives cancer to higher non-reversible hierarchical levels.

### Phase Analysis

Because the phase images correlate with the topographical ones, the signal’s driving frequency is associated with a phase shift, owing to adhesion, stiffness, or friction. Therefore, the standard statistical phase parameters of stained CRC histological sections AFM images were calculated. As for the AFM amplitude imaging, the *z*-height distribution of non-metastatic tissues has a broader dispersion around the mean value; Supplementary Fig. S6a online. Also, the RMS roughness values of metastatic tissues are lesser than for non-metastatic ones; Supplementary Fig. S6b online. Again, there is no clear distinction between metastatic and non-metastatic tissues. Rescaled range analysis is also applied for phase images: the Hurst exponent, Supplementary Figs. S6c,6d online, and the Hurst exponent distribution were extracted between the phase lines, Supplementary Fig. S6e online.

As for the amplitude images of the Hurst exponent, the differentiation between metastatic and non-metastatic tissues is unclear. However, surface and rescaled analysis bear noticeable similarities despite limiting metastatic information.

### Monofractal Image Analysis

Monofractal dimensionality *D_f_* of metastatic and non-metastatic tissues was calculated by cube counting, triangulation, power spectrum, and partition algorithms, Supplementary Fig. S7 online. The cube counting and the triangulation methods, Supplementary Figs. S7a,7d online provide a lower *D_f_* number for metastatic tissues than the other two methods, Supplementary Figs. S7b,7c online, where the fractal dimensionality of non-metastatic tissues is relatively more minor than that of metastatic ones. Overall, the four algorithms have no clear differentiation between metastatic and non-metastatic tissues.

## Discussion

Cancer is a multivariate and complex disease, and despite intense research starting as early as the last century, still, it represents a challenging issue. There are many reasons for it. Over the years, clinical methods applied favourable average practices. Nevertheless, cancer is highly heterogeneous even within the same cell and similar class. Consequently, an overall positive average outcome does not translate to individual positive results.

Moreover, despite the significant effort and the enormous resources devoted to cancer research, it is still unknown why drugs are more effective for some individuals and not for others. Besides other critical issues, deciphering cancer growth, metastatic progression and migration at the nanoscale is vital for survival. Likewise, metastasis shapes one of the six crucial hallmarks of cancer^2,65^; the others have been sustaining proliferating signalling, evading growth, suppression and activating invasion.

The intricacy is further increasing because almost 12 years after the seminal paper of Hanahan and Weinberg^2^, four additional cancer hallmarks highlight the disease’s complexity, signalling that traditional approaches need new strings in their bows^66^. First, it is now well understood that a novel interdisciplinary approach to the cancer menace is required, where biology, physics and mathematics, in an integrating step, could illuminate the dark pathways of cancer progression or even discover hidden physical laws of the phase transition between healthy and carcinogenic cells. Second, even if critical sporadic and uncorrelated contributions to cancer research were made from different physics and cell biophysics fields, their integration is still intermittent in cancer research. Third, the metastatic phase usually is clinically validated by biomarkers. So thought, even when the diagnosed metastatic phase is discovered with an optical histological examination of spatial resolution less than 500 *μm*, it represents a late and fatal stage. Fourth, the multistep process of invasion and metastasis mimics, under certain circumstances, a developmental program referred to as the Epithelial-Mesenchymal Transition (EMT)^67^. The basic idea of the current work is to identify, in the primary tumour sites, carcinoma cells with early morphological changes, which can indicate the activation of the EMT program. Indeed, during EMT, the carcinoma cells lose their cell-cell junctions and move apart, generating tiny but significant histological and cytological changes detected only at the nanoscale level with AFM.

The sill variogram values of metastatic CRC histological tissues from three patients are below 0.566, the threshold line for image resolutions 512 *px x* 512 *px* and *σ*= 10 *px* (1 *μm*). For different configurations of image resolutions and *σ* values, the metastatic threshold line could be adjusted accordingly. The metastatic threshold line from variograms between the metastatic and non-metastatic phases defines the borderline between death and extended survival of patients. Importantly, in the case of ambiguity, as for the nm2.4 tissue, higher moments than 2^nd^ order variograms remove any possible mixing between metastatic and non-metastatic tissues, Fig. 5, Supplementary Figs. S3,S4 online. Contrary to variograms and *θ*-statistics, P-value statistics verified that the rescaled range, surface, phase, and monofractal analysis does not distinguish between metastatic and non-metastatic tissues, and the correlation between metastasis and tissue mono-fractality is vague, Supplementary Tables T1,T2,T3 online. Indeed, during the transformation of single premalignant cells into cancerous^62^, the fractal dimensionalities do not necessarily imply the existence of fractal geometrical features.

In contrast, by applying P-value statistics in second-moment variograms and the null hypothesis that the mean sill values of metastatic and non-metastatic tissues are the same, the differentiation between metastatic and non-metastatic tissues (P-value) is confident with a probability of 99.99999%. The differentiating confidence for higher than 2^nd^ order variogram moments for metastatic and non-metastatic tissues is further improved. High order variograms of Gaussian residual filtering distinguish metastatic and non-metastatic tissues by categorizing a well-defined threshold. The reason is that Gaussian filtering differentiates the *z*-heights features with size less than 97.5, 194, and 388 *nm* (for image resolutions 512 *px x* 512 *px*, 256 *px x* 256 *px*, 128 *px x* 128 *px*, respectively). This result agrees with previous work^62^, where microvilli, micro ridges, and glycocalyx are responsible for the pericellular brush surface geometry structure. AFM imaging includes information from the cell’s surface, random cell volume cross-sections, CRC histological tissue encloses and ECM. Therefore, Gaussian filtering differentiates small biological features between metastatic and non-metastatic phases in the CRC-ECM system.

Besides the practical utility of variograms in cancer prognosis, grouping the well-defined threshold sill lines for metastatic and non-metastatic CRC tissues have broader implications in cancer research. Undeniably, the differentiation between the metastatic and non-metastatic phases defines two different hierarchies in the CRC cell-ECM system. Generally, dynamical systems, such as cancerous ones, have structural (hardware) and functional (software) connotations that form ensembles of successfully interacting nested sets and subunits of variables and parameters. Also, as the complexity of structural and functional systems depends on the number of their components and interconnections, it is inversely proportional to the stability and the degrees of freedom. It thus defines a particular hierarchical state (level). Furthermore, the systems afford a specific state-space-time description with certain collective properties (e.g., statistical moments, convolutions, distribution functions). From that state, during the evolution process across the dynamical paths, the systems within “limited-time series” are commonly driven to lower complexities with fewer degrees of freedom and thus to more stable states (high viability).

Therefore, the dynamical systems evolve from lower hierarchical levels with many degrees of freedom and high complexities to higher hierarchical levels with fewer degrees of freedom and lower complexities. Besides structural hierarchies, the systems are characterized by the dynamic of formation. The higher levels receive selective information from the lower levels through the cognition (memory)^68^ of collective properties. In turn, they exercise negative feedback control commands on the dynamics of the lower levels in their effort to occupy successfully higher hierarchical levels. Therefore, interactive systems are characterized by mutual “simulation”. One dynamic system, say a non-metastatic one, tries to simulate another with fewer degrees of freedom and higher stability (metastatic system). Thus, a non-metastatic system will eventually occupy higher hierarchical levels of lower complexity with higher stability. The opposite route, the evolution from higher hierarchical levels to lower ones, requires the expenditure of additional information energy (entropy). Therefore, in most cases, the reverse process is energetically unfavoured. Along the above lines, the selective differentiation between metastatic and non-metastatic groups evinces the dynamic evolution of different hierarchical carcinogenic states during the stages of disease progression. The advancement ranges from lower carcinogenic hierarchical levels of higher complexity and low stability (premalignant conditions) to higher ones that are less complex and stable (metastatic forms). The evolutionary dynamic might well explain the heterogeneous chemotherapy results. If the hierarchical dynamic is deciphered, the cancer therapeutic protocols and road map might change.

## Materials and methods

### Histological Tissue Preparation

CRC human histological tissues were prepared at the University Hospital “Federico II” in Naples, Italy, using anonymous numerical codes. Human tissues were handled and prepared following the Helsinki protocol (http://www.wma.net/en/30publications/10policies/b3/) and the practices approved by the legal guardians of Comitato Etico, Universita Frederico II (protocol article 152-18/ 13/06/2018) and the Bioethics Committee of NHRF (reference number 2/13-04-2022). The tissue samples were labelled 1) according to the tumour site (right colon, transverse, left colon, rectosigmoid), 2) the pathological classification (Cancer Control UICC, 2017, T, N, M), 3) the vascular hematic, vascular lymphatic and perineural invasion, and 4) the surgical resection margin status. Necrosis, neoplastic cellular percentage, desmoplasia, and tumour-infiltrating lymphocytes were assessed by optical microscopy. The mucinous acellular component was categorized as absent (<1%) and present (≤50% or >50%).

The tumour histological sections were collected on glass slides in FFPE blocks. Before AFM imaging, they dewaxed at 60°*C*. Then, the wash was for 300 *s* in three steps with xylene, and xylene traces were removed by three washing steps in 100 % ethanol for 300 *s* each time. After that, slides were further washed in 95% ethanol for 300 *s* and once in distilled water again for 300 *s*. Samples were stained with hematoxylin solution, according to Mayer (Sigma Aldrich Chemie GmbH, Riedstr. 2-D89555 Steinheim49 7329 970, 1.044 *grml*^-1^ at 20° *C*) and dried in the air for about 600 *s* at 20°*C*.

### AFM Image Analysis

Sixteen metastatic/non-metastatic fixed histological tissues were imaged by Innova AFM (Bruker/Veeco, Inc., Santa Barbara, CA) operating in tapping mode with phosphorus (n)-doped silicon cantilever (RTESPA, Bruker, Madison, 120 WI, USA) with a nominal tip diameter of 8-10 *nm*, and nominal spring constant of 40 *N/m* at 300 *kHz* resonance frequency.

Surface image quality was optimized by lowering the scan rate to 0.2 *Hz*. All images were acquired with 50 *μm x* 50 *μm* scan sizes, 512 *x* 512 data point resolution, and pixel size 97.656 *nm*. Besides height, amplitude and phase images were also recorded. The AFM was installed on a vibration isolation table (minus k technology BM-10) to compensate for regular environmental vibrations and placed inside an acoustic enclosure (Ambios technologies Isochamber) for thermal and building vibrations isolation. The AFM imaging was performed in air at constant ambient temperature.

### Histological Tissue Optical Analysis

Before AFM imaging, optical microscopy was used for metastatic identification. First, the paraffin-stained CRC histological sections were placed under a transmitted light optical microscope (Carl Zeiss, Primovert microscope) with magnifications 4*x*, 20*x*, and 40*x*. Then, the AFM probe was positioned in the identified image areas.

### Gaussian Filtering Residuals RMS Deviation

A *3D* Gaussian filter is applied to the original image for each AFM image. The Gaussian cubic filter size (kernel) was set to 31 *px*, a standard deviation *σ* of 5 *px* in every dimension. The residuals of the Gaussian filter (a high-pass filter that represents the small scale roughness of the surface) consist primarily of the spatial frequencies below the cut-off wavelength (6*σ+1* = 31 *px* or ~ 3 *μm*, 1 *px* =97.65 *nm* for 512 *px x* 512 *px* image resolution), with some leakage of higher spatial frequencies. The statistical measure of the height differences for all possible point pairs of an area at a particular scale, the RMS deviation *D*(*h*) is determined for each lag vector h = (±*v*, ±*p*), then scaled with lag vectors’ magnitude.

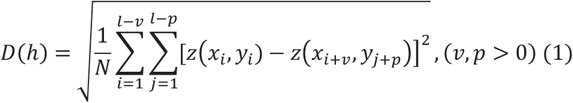

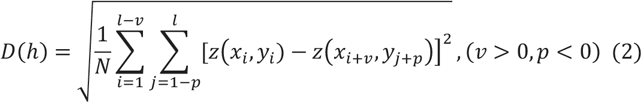

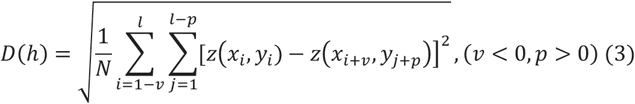

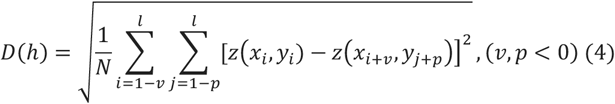

where *l* stands for the size of the image and *N* is the number of sample points separated by 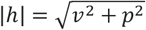.

The RMS deviation as a function of lag vectors in all directions is depicted in *2D* or *1D* plots (variograms).

*1D* plots depict the RMS deviation between all points spaced apart by 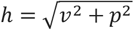 Alternatively called empirical or experimental variograms/semivariograms. The empirical variograms were calculated as the average of the square differences of the values *z*(*x_i_*, *y_i_*), *z*(*x_i+v_*, *y_j+p_*) of all pairs of locations that fall within length intervals, *h* (lags).

The sill value in variograms depicts zero correlation of lag vectors, visualized with variograms’ flattening off, Fig. 4a. The analysis was made for three different image resolutions, 512, 256 and 128 *px* per axis and three different Gaussian filtering standard deviation values of 2.5, 5 and 10 *px*.

### Moments of Gaussian Filtering Residuals Variogram

Various Gaussian Filtering Residuals Variogram moments were calculated as an extension of the previous method. For q = [0.5,1,2,3,4, 5], the generalised variogram *γ*(*h, q*) was evaluated.

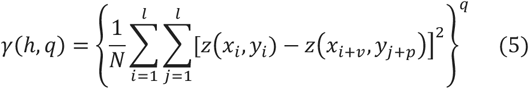

Then the generalised variogram sill was calculated and compared for metastatic/non-metastatic samples. The small order moments, 0 < *q* < 2, are responsible for the core of the probability density function (PDF), while higher moments contribute to the tails of the PDF. For *q* = 1 generalised variogram is the empirical variogram. Comparing generalised and simple variogram sills of metastatic/non-metastatic samples leads to clear differentiation of samples.

### Theta Statistics

Inclination-slope distributions were applied for metastatic and non-metastatic AFM images using the Gwyddion software^69^. The polar angle *θ* between the horizontal plane and the “central derivative plane” in every pixel is related to the surface-profile gradient 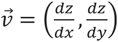 via the equation 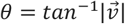. The polar angle *θ* is always positive and rises with a slope 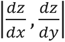 while the integral ∫ *ρ*(*θ*)*dθ*, for 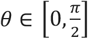 is normalized to one and 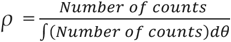. For quantified comparison between different slope distributions the skewness 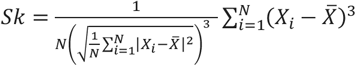 and kyrtosis 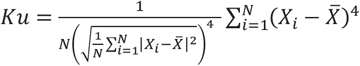 of slope distribution is calculated^63^.

### Rescaled Range Analysis

The Hurst exponent of every line of AFM image was calculated using the rescaled range analysis of Hurst^64^. The algorithm for these calculations was designed and run in MATLAB. 9.4.0.813654 (R2018a), The MathWorks Inc.; Natick, MA, the USA based on Weron’s algorithm^70^. Then, the mean value of the Hurst exponent of every AFM image was calculated, and the histogram was plotted. In addition, the *2D* AFM image was transformed into a *1D* array by putting every line after another. Finally, the Hurst exponent of every *1D* AFM image was calculated with the same methodology. The same analysis was also performed on “phase” images.

### Surface Statistics

Several different parameters were used for the first qualitative evaluation of surface characteristics of metastatic and non-metastatic samples. First, was calculated the average *z*-height (*nm*), an arithmetic mean defined as the sum of all height values divided by the number of data points 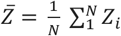. Next, the RMS roughness was calculated, which is the square root of the mean value of the squares of the distance of the points from the image mean value, 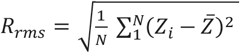.

### Phase Spectra

The AFM tapping mode generates the phase images, and the signal’s frequency (phase) is a function of the driving frequency adjusted to be at the actual probe resonance, as it is shifted due to the tip-sample forces. Tip height variation of phase images correlates with the topographical ones. The signal’s driving frequency is associated with a phase shift, owing to adhesion, stiffness, or friction, and the interaction between tip and surface will cause the lag oscillation. The RMS phase roughness, mean phase, mean phase Hurst exponent of 512 lines, and Hurst exponent of *1D* image vector was calculated for all phase AFM images.

Statistical analysis was applied to both images. The average phase shift *(V)* is an arithmetic mean defined as the sum of all height values divided by the number of data points 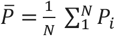. The RMS roughness is the square root of the mean value of the squares of the distance of the points from the image mean value, 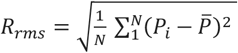.

### Monofractal Image Analysis

The self-affine properties in a specific range of scales were analyzed using monofractal analysis. Four methods calculated the fractal dimension *D_f_* using “Gwyddion, SPM data visualization and analysis tool.” The methods used to calculate *D_f_* are cube counting, triangulation, and partition. Cube counting arises from the definition of box-counting fractal dimension where an *l* cubic lattice constant superimposes on the *z*-expanded surface. First, *l* is set at *X/2* (*X* is the surface’s edge length), providing a lattice of 2^3^ cubes *N(l)*, containing at least one pixel. The lattice constant *l* is reduced stepwise by a factor of two, and the process repeats until *l* equals the distance between two adjacent pixels. The slope of a plot of *log(N(l))* versus *log(1/l)* gives *D_f_*. Triangulation is similar to the previous method. A grid of unit dimension *l* is placed on the surface, defining the location of several triangle vertices. For *l* = *X/4*, 32 triangles of different areas inclined at different angles with the *xy* plane cover the surface. The areas of all triangles are calculated and summed to approximate the surface area *S(l)* for a given *l*. Next, the grid size decreases by a successive factor of 2, and the process continues until *l* equals the distance between two adjacent pixel points. The slope of a plot of *log(S(l))* versus *log(1/l)* is the number *D_f_* - 2. The partitioning algorithm is based on the scale dependence of the variance of fractional Brownian motion. One divides the entire surface into equal-sized squared boxes, and the variance (power of RMS heights) was calculated for the particular box size. The slope value *β* of a least-square regression line fits the data points in the log-log plot of variance extracts *D_f_’s* as *D_f_* = 3 − *β/2*.

## Author contributions

VG, AF, ACC, ES conceive the idea and design the work. VG, AF, ZK, ES perform and analyze the AFM and optical images. FP, UM, CDL, GT prepared the histological CRC sections. ACC, ES conceive the idea of using variograms in AFM cancer histological sections. VG calculates the moments of residual Gaussian filtering, theta analysis, surface statistics, rescaled, phase, and fractal analysis. ACC conceive the idea of hierarchical levels and non-reversible dynamics in cancer progression.

## Data availability statement

All data published in this article are available upon request.

