## supplemental material online for "Nanoscale prognosis of colorectal cancer metastasis from AFM image processing of histological sections"

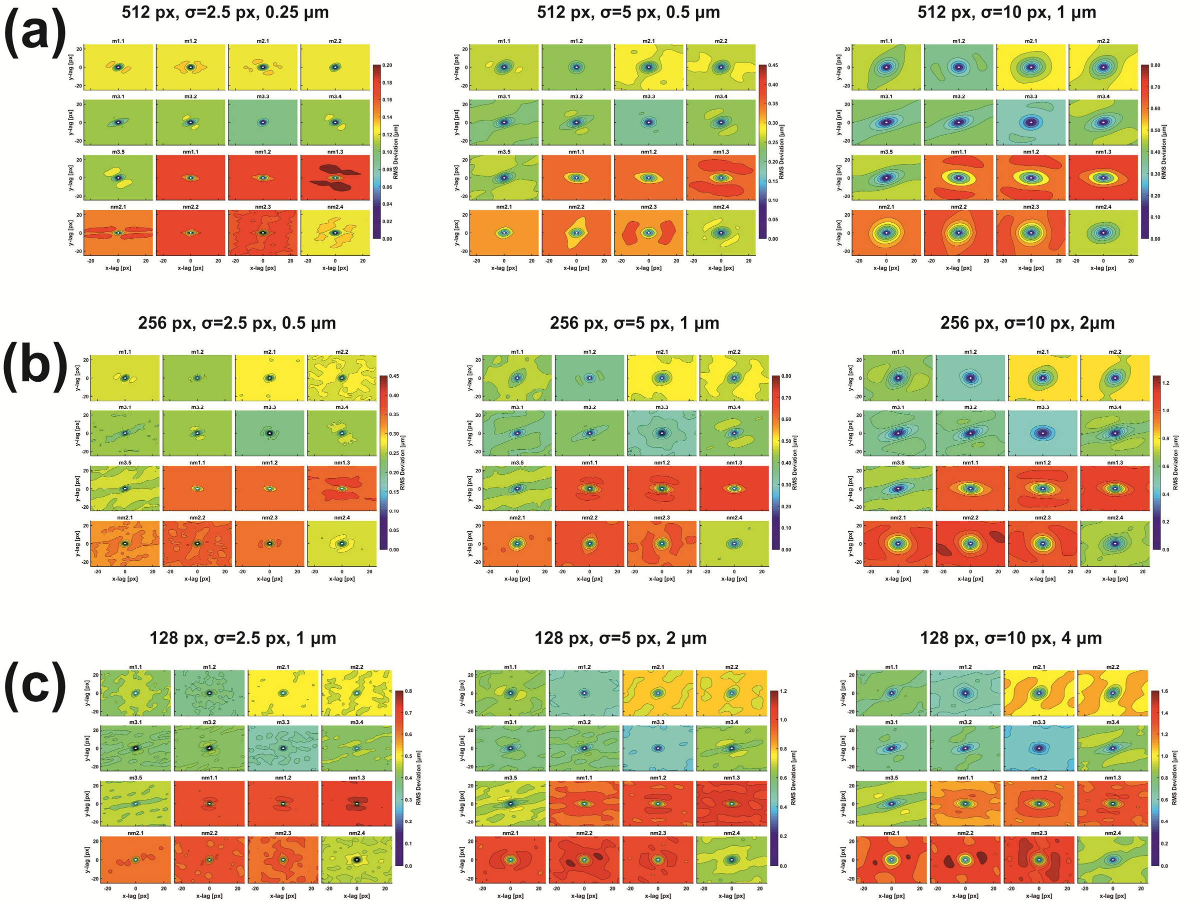


**Fig. S1 *2D* variograms of the residuals of the Gaussian filtered AFM images of metastatic and non-metastatic histological tissues for (a)** AFM image resolution 512 *px* x 512 *px,* and *σ* 2.5, 5 and 10 *px*. **(b)** AFM image resolution 256 *px* x 256 *px*, and *σ* 2.5, 5 and 10 *px*. **(c)** AFM image resolution 128 *px* x 128 *px* and *σ* 2.5, 5 and 10 *px*. The magnitude of RMS deviation of closed contour areas diverges for metastatic and non-metastatic phases except for sample nm2.4, which shows metastatic behaviour. The sample nm2.4 attains the correct metastatic state for higher moments ($q$>2).

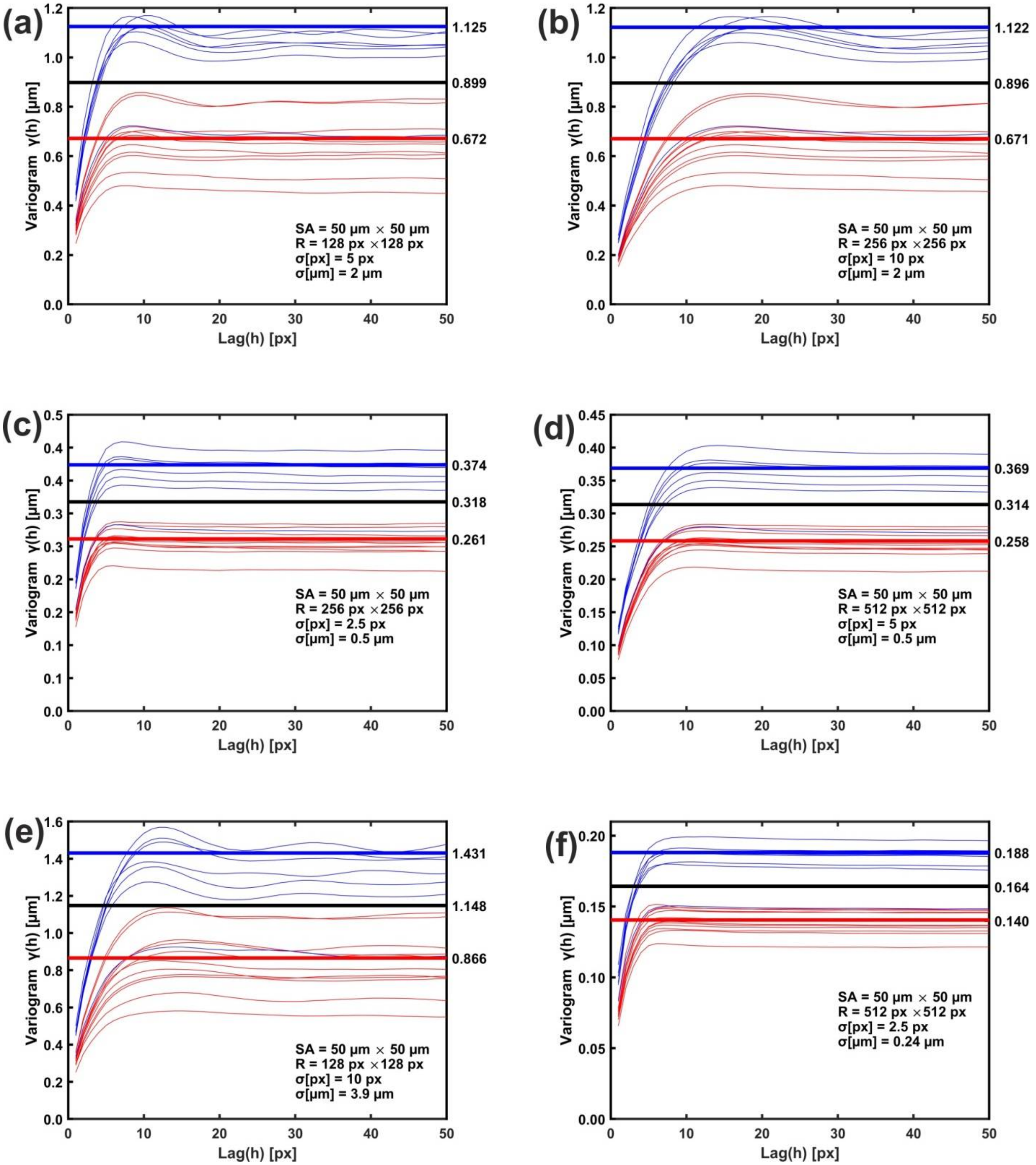


**Fig. S2 Non-metastatic (blue lines) and metastatic (red lines) *1D* variograms Variograms of low-resolution images and large *σ's* bear wider gaps and high uncertainty between the mean sill values of metastatic and non-metastatic variogram bands. (a,b)** The pair 128 *px x* 128 *px*, *σ* 5 *px (2 µm)* and 256 *px x* 256 *px*, *σ* 10 *px* (2 *µm*) hold the same metastatic threshold values. **(c,d)** The pair 256 *px x* 256 *px*, *σ* 2.5 *px (0.5 µm*) and 512 *px x* 512 *px*, *σ* 5 *px (0.5 µm*), hold the same metastatic threshold values. **(e,f)** The pair 128 *px x* 128 *px, σ* 10 *px (3.9 µm)* and 512 *px x* 512 *px, σ* 2.5 *px* (0.24 *µm*) hold the same metastatic threshold values.

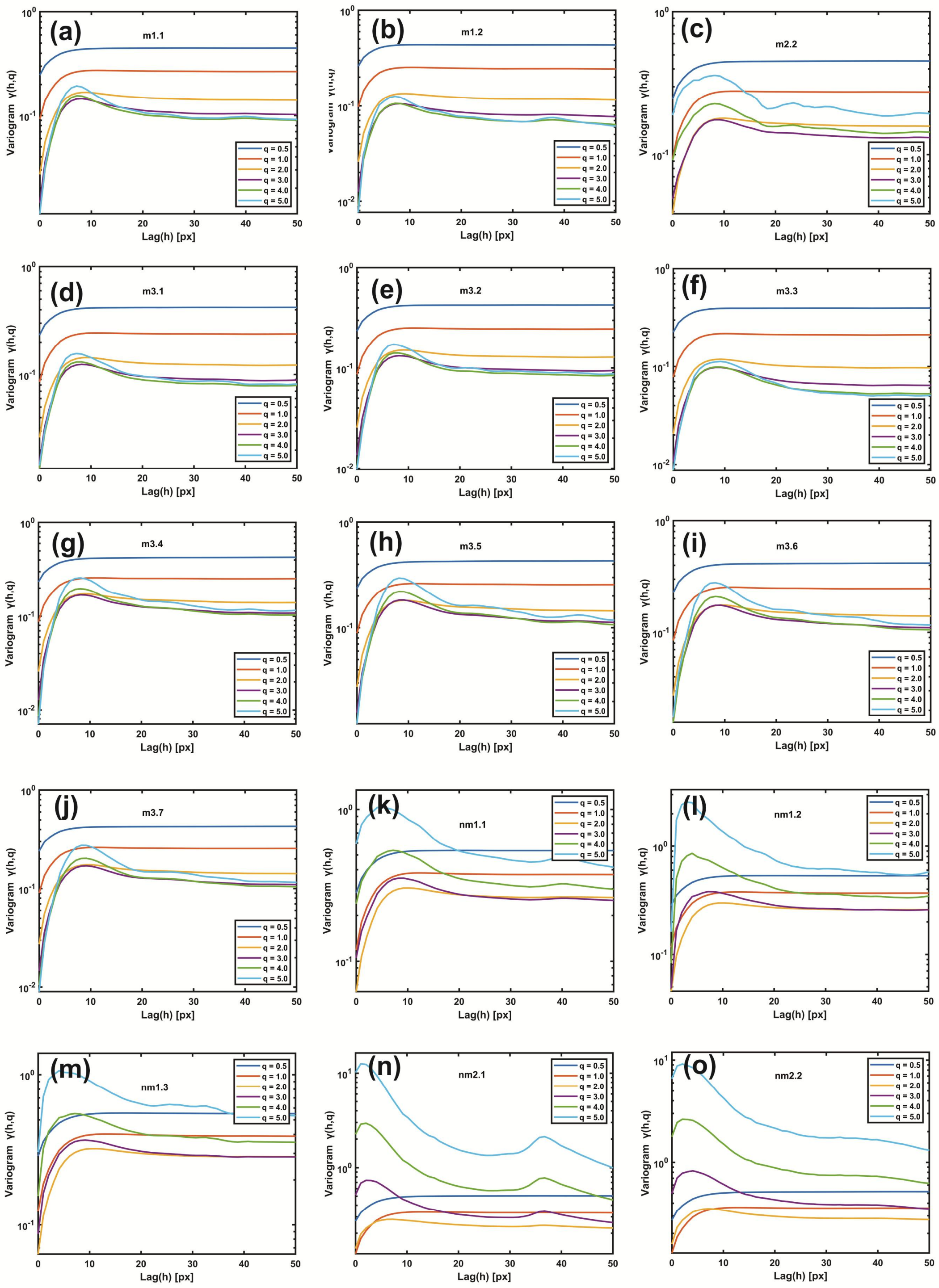


**Fig. S3 Gaussian filtering residuals variograms of different moments *(q).*** **(a)-(j)** for metastatic tissues. **(k-o)** for non-metastatic tissues.

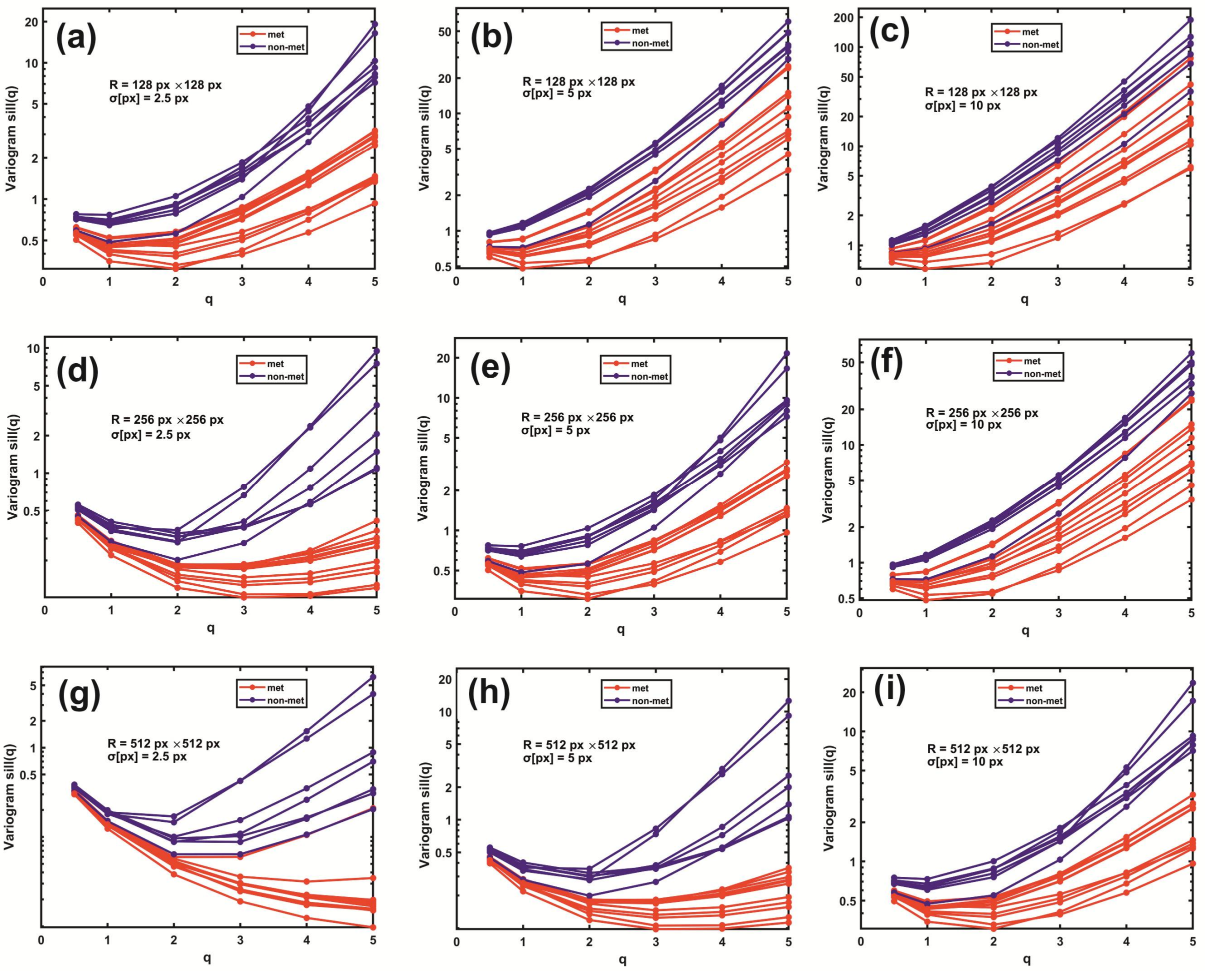


**Fig. S4 Variogram sill values for different image resolutions and Gaussian filtering *σ* *vs.* scaling exponents *q* for metastatic (red) and non-metastatic (blue) tissues. (a-c)** 128 *px x* 128 *px*, *σ* 2.5, 5 *px* and 10 *px*. **(d-f)** 256 *px x* 256 *px*, *σ* 2.5, 5 and 10 *px*. **(g-i)** 512 *px x* 512 *px*, *σ* 2.5, 5 *px* and 10 *px*. The nm2.4 tissue, the non-successful sample, in the *1D* variograms metastatic threshold criterion, performs as metastatic for higher moments (*q*>2), adopting the correct non-metastatic state in agreement with the subjective optical microscopic examination. The differentiation between metastatic and non-metastatic tissues is improved for high *q* values.

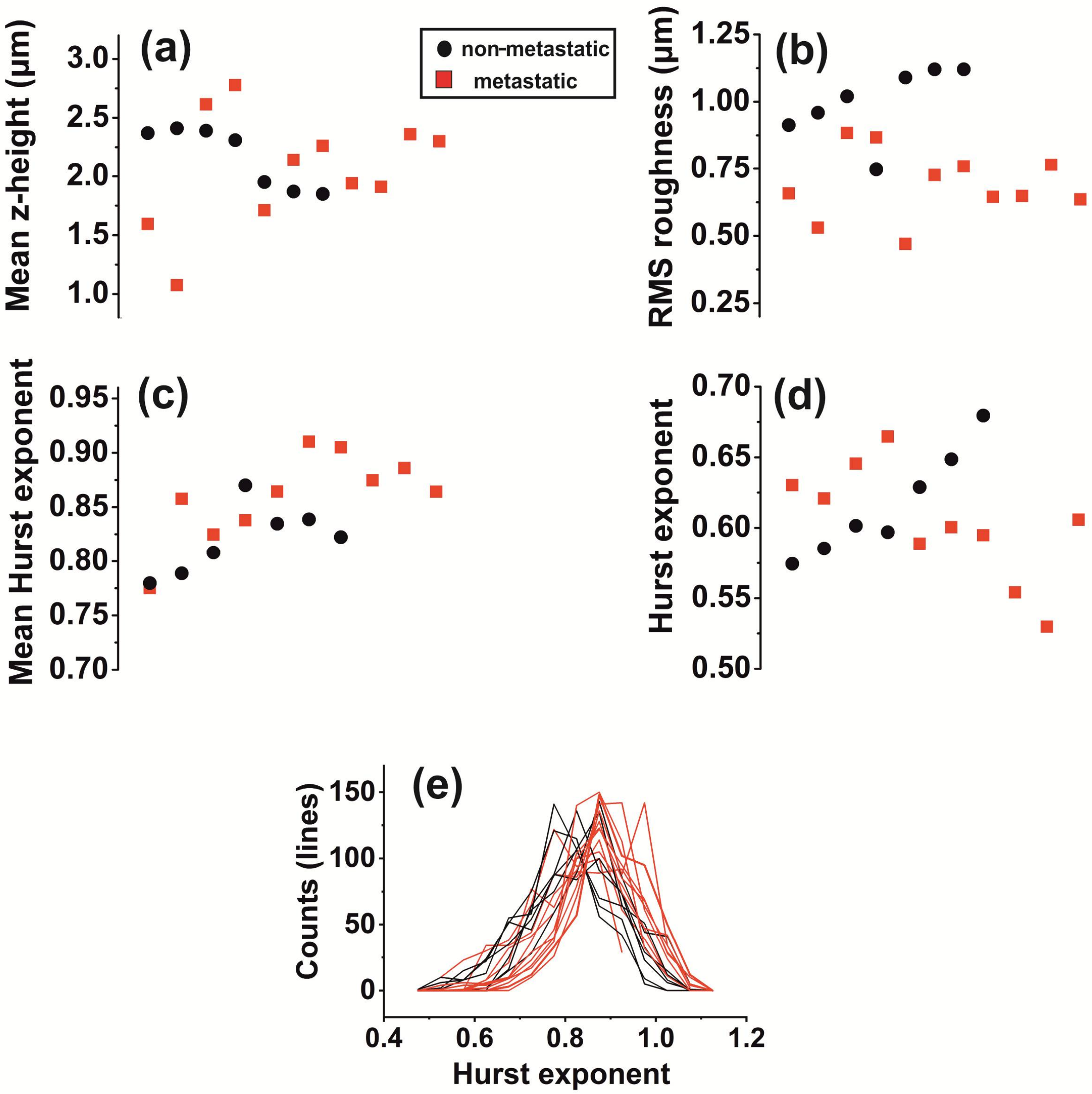


**Fig. S5 Standard surface statistical parameters and rescale range analysis/surface statistics of AFM images of CRC histological sections. (a)** Mean *z*-height distribution. **(b)** RMS roughness. **(c)** Mean Hurst exponent. **(d)** Hurst exponent. **(e)** Hurst exponent distribution. The differentiation between metastatic and non-metastatic sections is unclear.

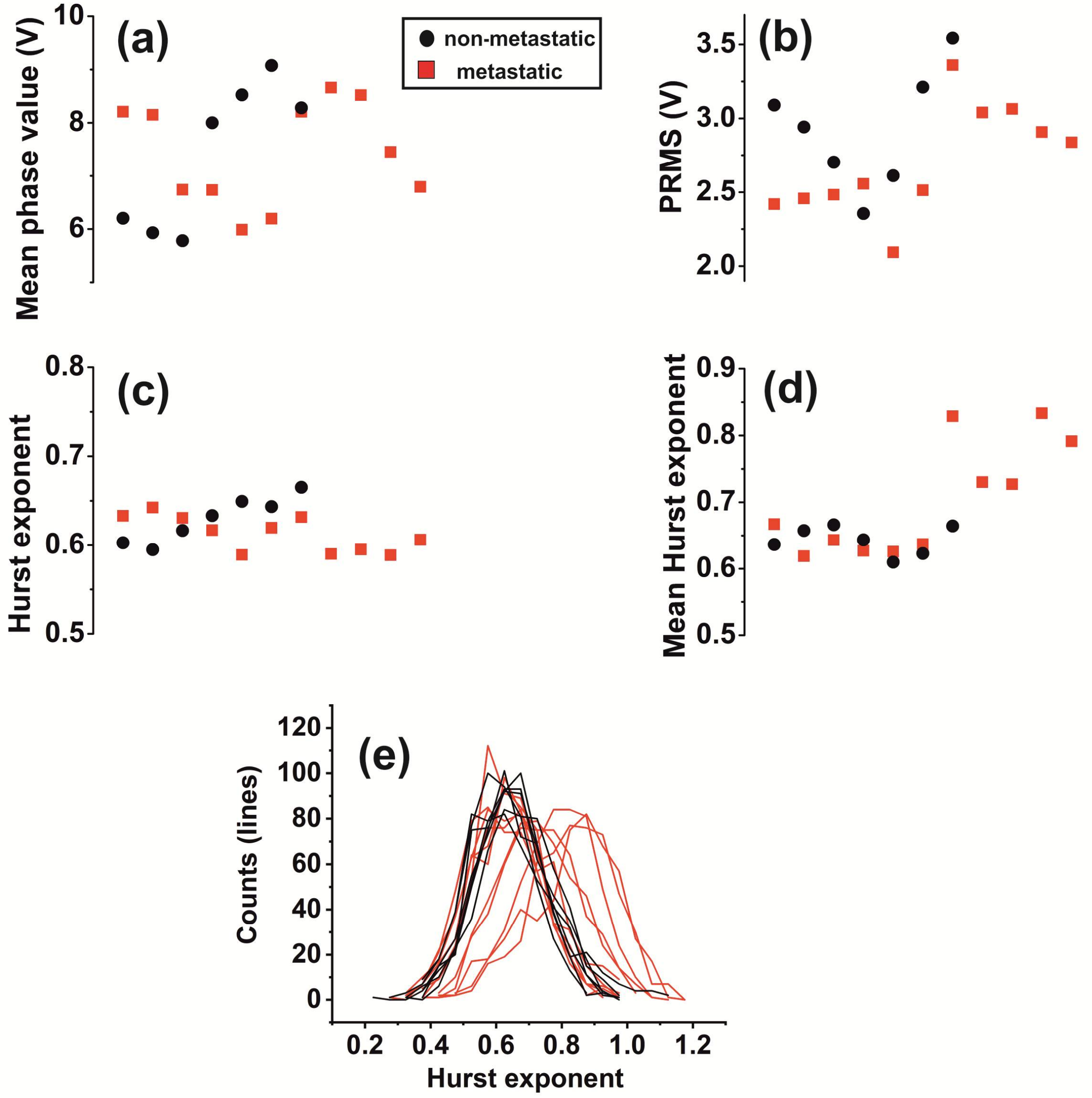


**Fig. S6 Surface statistical phase spectra of CRC metastatic (red squares) and non-metastatic (black circles) tissue AFM images. (a)** Mean phase value. **(b)** Phase RMS roughness. **(c)** Hurst exponent. **(d)** Mean Hurst exponent. **(e)** Hurst exponent distribution. The differentiation between metastatic and non-metastatic sections is unclear.

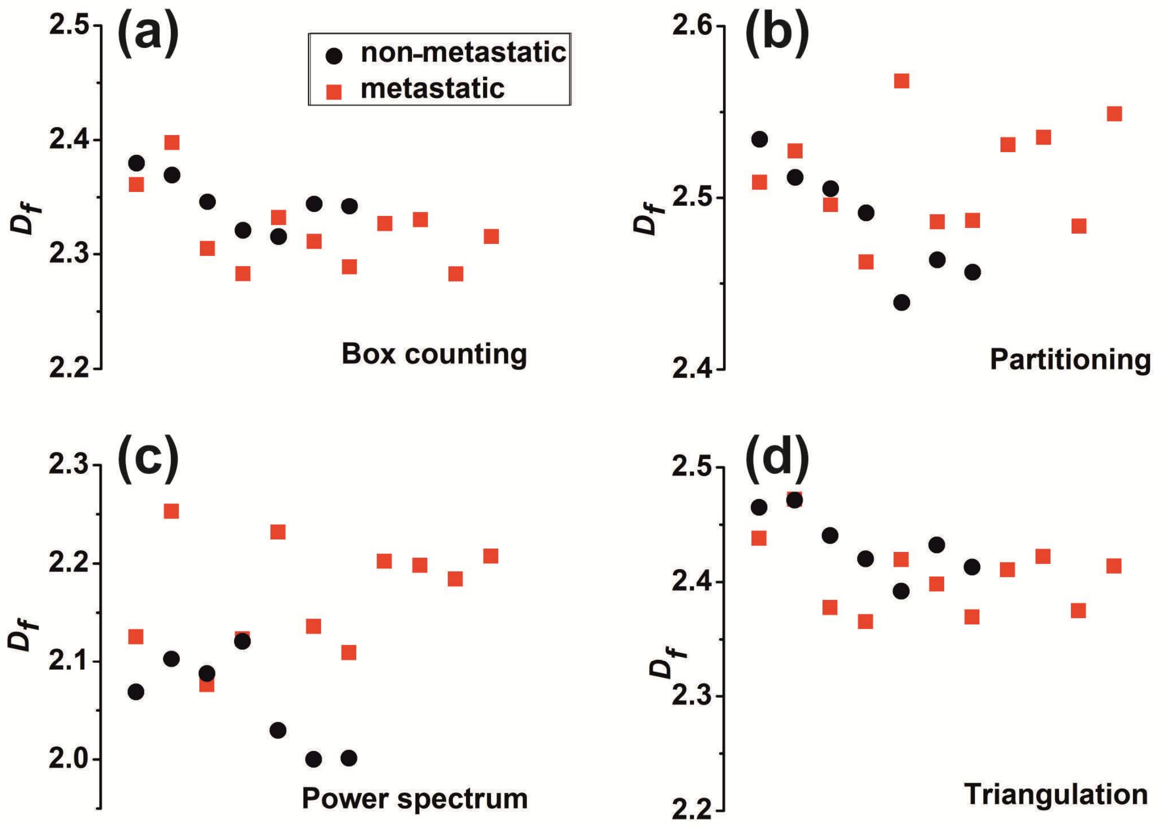


**Fig. S7 Fractal dimension $D_f$ of images of CRC metastatic (red squares) and non-metastatic (black circles) tissue AFM images calculated with four different methods. (a)** Box counting. **(b)** Partitioning. **(c)** Power spectra. **(d)** Triangulation. The differentiation between metastatic and non-metastatic sections is unclear.

**Two-sample t-test analysis**

The independent two-sample t-test analysis inspects whether or not the means of two independent samples are equal or whether they differ by a pre-defined value and creates a confidence interval for the difference of the sample means. A two-sample t-test analysis was performed, taking the 11 variogram sills of metastatic (M) and 7 variogram sills of non-metastatic (NM) tissues as independent samples. The null hypothesis is that the variogram mean sill values of M and NM samples are equal. Calculations of the *t*-test statistic and p-value determine whether or not to reject the null hypothesis. A smaller p-value means that there is strong evidence in favour of the alternative hypothesis (rejection). Considering that the proposed variogram sill threshold for M or NM characterization is the median value between mean values of M/NM samples variogram sills, the confidence interval for the difference provided by t-test is the confidence interval of the threshold value.

The null hypothesis takes the form $\mu_{NM} - \mu_{M} = 0$, while the alternative hypothesis is $\mu_{NM} - \mu_{M} \neq 0$. A two-tailed t-test with non-equal variances was performed. An approximate test statistic, *t* is used:

$$t = \frac{\mu_{NM} - \mu_{M}}{\sqrt{\frac{s_{NM}^2}{N_{NM}} + \frac{s_{M}^2}{N_{M}}}}$$

where $\mu$ and $s^2$ are the means and the variances of the NM and M populations, respectively. A t-distribution with $v$ degrees of freedom is used to approximate the distribution of *t* where

$$v = \frac{\left(\frac{s_{NM}^2}{N_{NM}} + \frac{s_{M}^2}{N_{M}}\right)^2}{\frac{\left(\frac{s_{NM}^2}{N_{NM}}\right)}{N_{NM} - 1} + \frac{\left(\frac{s_{M}^2}{N_{M}}\right)}{N_{M} - 1}}$$

Then the $t$ value is compared to the critical value, and the null hypothesis is rejected if $|t| > t_{\sigma/2}$ where $t_{\sigma/2}$ is the critical value of the t-distribution with $v$ degrees of freedom and $\sigma$ significance level. The p-value was also compared with $\sigma$ significance level, 0.05 in our case. The upper and lower $(1 - \sigma) \times 100\%$ confident limits for the mean difference $\mu_{NM} - \mu_{M}$ are calculated as:

$$\left[ (\mu_{NM} - \mu_{M}) - t_{\frac{\sigma}{2}}\sqrt{\frac{s_{NM}^2}{N_{NM}} + \frac{s_{M}^2}{N_{M}}}\ , (\mu_{NM} - \mu_{M}) + t_{\sigma/2}\sqrt{\frac{s_{NM}^2}{N_{NM}} + \frac{s_{M}^2}{N_{M}}} \right]$$

Data for the t-test analysis were taken from Supplementary Table T1, and the t-test results, the corresponding p-values and the threshold confidence intervals are shown in Supplementary Table T2, and Supplementary Table T3.

| Resolution | 512 *px* × 512 *px* | | | | | | 256 *px* × 256 *px* | | | | | | 128 *px* × 128 *px* | | | | | |
|---|---|---|---|---|---|---|---|---|---|---|---|---|---|---|---|---|---|---|
| Sigma [*px*] | σ=10 *px* | | σ=5 *px* | | σ=2.5 *px* | | σ=10 *px* | | σ=5 *px* | | σ=2.5 *px* | | σ=10 *px* | | σ=5 *px* | | σ=2.5 *px* | |
| Sigma [*μm*] | σ=1 *μm* | | σ=0.5 *μm* | | σ=0.24 *μm* | | σ=2 *μm* | | σ=1 *μm* | | σ=0.5 *μm* | | σ=3.9 *μm* | | σ=2 *μm* | | σ=1 *μm* | |
| Sample | Sill (*μm*) | Range (*px*) | Sill (*μm*) | Range (*px*) | Sill (*μm*) | Range (*px*) | Sill (*μm*) | Range (*px*) | Sill (*μm*) | Range (*px*) | Sill (*μm*) | Range (*px*) | Sill (*μm*) | Range (*px*) | Sill (*μm*) | Range (*px*) | Sill (*μm*) | Range (*px*) |
| **m1.1** | 0.467 | 21 | 0.274 | 12 | 0.149 | 7 | 0.699 | 16 | 0.469 | 11 | 0.277 | 6 | 0.852 | 12 | 0.699 | 8 | 0.472 | 6 |
| **m1.2** | 0.395 | 18 | 0.254 | 11 | 0.152 | 6 | 0.533 | 17 | 0.395 | 9 | 0.256 | 6 | 0.680 | 15 | 0.534 | 9 | 0.397 | 5 |
| **m2.1** | 0.515 | 23 | 0.284 | 13 | 0.150 | 8 | 0.854 | 19 | 0.519 | 12 | 0.287 | 7 | 1.136 | 13 | 0.858 | 10 | 0.526 | 6 |
| **m2.2** | 0.506 | 23 | 0.279 | 13 | 0.147 | 8 | 0.843 | 19 | 0.509 | 12 | 0.283 | 7 | 1.113 | 13 | 0.846 | 10 | 0.515 | 6 |
| **m3.1** | 0.410 | 20 | 0.245 | 12 | 0.135 | 7 | 0.603 | 17 | 0.412 | 10 | 0.248 | 6 | 0.765 | 15 | 0.603 | 9 | 0.414 | 5 |
| **m3.2** | 0.422 | 20 | 0.252 | 12 | 0.137 | 7 | 0.616 | 17 | 0.424 | 10 | 0.255 | 6 | 0.777 | 16 | 0.616 | 9 | 0.426 | 5 |
| **m3.3** | 0.349 | 19 | 0.219 | 11 | 0.124 | 7 | 0.481 | 15 | 0.350 | 10 | 0.221 | 6 | 0.581 | 15 | 0.481 | 8 | 0.352 | 5 |
| **m3.4** | 0.446 | 22 | 0.258 | 12 | 0.139 | 8 | 0.683 | 17 | 0.449 | 11 | 0.261 | 6 | 0.902 | 15 | 0.685 | 9 | 0.453 | 6 |
| **m3.5** | 0.465 | 22 | 0.263 | 12 | 0.140 | 8 | 0.719 | 17 | 0.467 | 11 | 0.266 | 6 | 0.949 | 16 | 0.719 | 9 | 0.471 | 6 |
| **m3.6** | 0.443 | 21 | 0.254 | 12 | 0.133 | 9 | 0.701 | 20 | 0.445 | 10 | 0.257 | 6 | 0.964 | 15 | 0.704 | 10 | 0.450 | 5 |
| **m3.7** | 0.441 | 20 | 0.262 | 12 | 0.142 | 7 | 0.646 | 17 | 0.443 | 10 | 0.265 | 6 | 0.806 | 13 | 0.646 | 8 | 0.446 | 5 |
| **Mean** | **0.442** | **20.8** | **0.258** | **12.0** | **0.140** | **7.5** | **0.671** | **17.4** | **0.444** | **10.5** | **0.261** | **6.2** | **0.866** | **14.4** | **0.672** | **9.0** | **0.447** | **5.5** |
| **nm1.1** | 0.692 | 20 | 0.381 | 13 | 0.191 | 9 | 1.061 | 17 | 0.696 | 10 | 0.386 | 7 | 1.275 | 11 | 1.063 | 9 | 0.701 | 5 |
| **nm1.2** | 0.697 | 21 | 0.377 | 14 | 0.188 | 10 | 1.100 | 17 | 0.701 | 11 | 0.383 | 7 | 1.356 | 12 | 1.104 | 9 | 0.708 | 6 |
| **nm1.3** | 0.757 | 21 | 0.403 | 14 | 0.199 | 13 | 1.165 | 16 | 0.761 | 10 | 0.409 | 7 | 1.383 | 11 | 1.167 | 8 | 0.767 | 5 |
| **nm2.1** | 0.633 | 24 | 0.339 | 14 | 0.180 | 8 | 1.123 | 20 | 0.637 | 12 | 0.343 | 7 | 1.511 | 12 | 1.128 | 10 | 0.645 | 7 |
| **nm2.2** | 0.670 | 22 | 0.362 | 14 | 0.188 | 8 | 1.165 | 21 | 0.674 | 12 | 0.367 | 7 | 1.570 | 12 | 1.169 | 11 | 0.681 | 6 |
| **nm2.3** | 0.651 | 22 | 0.350 | 13 | 0.182 | 11 | 1.118 | 19 | 0.656 | 12 | 0.356 | 7 | 1.490 | 13 | 1.120 | 10 | 0.662 | 6 |
| **nm2.4** | 0.482 | 20 | 0.280 | 13 | 0.150 | 8 | 0.722 | 17 | 0.484 | 10 | 0.283 | 6 | 0.925 | 14 | 0.723 | 8 | 0.486 | 5 |
| **Mean** | 0.655 | 21.4 | 0.356 | 13.6 | 0.183 | 9.6 | 1.065 | 18.1 | 0.658 | 11.0 | 0.361 | 6.9 | 1.359 | 12.1 | 1.068 | 9.3 | 0.664 | 5.7 |
| **Mean (-nm2.4)** | **0.683** | **21.7** | **0.369** | **13.7** | **0.188** | **9.8** | **1.122** | **18.3** | **0.687** | **11.2** | **0.374** | **7.0** | **1.431** | **11.8** | **1.125** | **9.5** | **0.694** | **5.8** |
| **Threshold** | 0.548 | | 0.307 | | 0.162 | | 0.868 | | 0.551 | | 0.311 | | 1.112 | | 0.870 | | 0.556 | |
| **Threshold (-nm2.4)** | **0.563** | | **0.314** | | **0.164** | | **0.896** | | **0.566** | | **0.318** | | **1.148** | | **0.899** | | **0.571** | |

**Supplementary Table T1.** Variogram parameters (sill, range), mean value and threshold, the arithmetic mean of sill values of metastatic (M) and non-metastatic (NM) tissues, with and without the nm2.4 sample (red colour), for different AFM image resolution and standard deviations $\sigma$ in *px* and $\mu m$.

| Resolution ($px$ x $px$) | $\sigma$ ($px$) | Tissue | Mean ($\mu m$) | StDev ($\mu m$) | SE Mean ($\mu m$) | D ($\mu m$) | StDev of D ($\mu m$) | 95% CI for D ($\mu m$) | Threshold ± ( 95% CI for D) ($\mu m$) | NM ≠ M: P-Value |
|---|---|---|---|---|---|---|---|---|---|---|
| **512 x 512** | **10** | **NM** | 0.683 | 0.043 | 0.018 | 0.242 | 0.046 | 0.050 | **0.563±0.050** | 3.14E-07 |
| | | **M** | 0.442 | 0.048 | 0.014 | | | | | |
| | **5** | **NM** | 0.369 | 0.023 | 0.009 | 0.110 | 0.020 | 0.025 | **0.314±0.025** | 5.60E-06 |
| | | **M** | 0.258 | 0.018 | 0.005 | | | | | |
| | **2.5** | **NM** | 0.188 | 0.007 | 0.003 | 0.048 | 0.008 | 0.008 | **0.164±0.008** | 2.53E-08 |
| | | **M** | 0.140 | 0.008 | 0.003 | | | | | |
| **256 x 256** | **10** | **NM** | 1.122 | 0.040 | 0.016 | 0.451 | 0.096 | 0.082 | **0.896±0.082** | 1.49E-08 |
| | | **M** | 0.671 | 0.114 | 0.034 | | | | | |
| | **5** | **NM** | 0.687 | 0.043 | 0.018 | 0.243 | 0.047 | 0.050 | **0.566±0.050** | 2.62E-07 |
| | | **M** | 0.444 | 0.049 | 0.015 | | | | | |
| | **2.5** | **NM** | 0.374 | 0.024 | 0.010 | 0.112 | 0.020 | 0.026 | **0.318±0.026** | 5.64E-06 |
| | | **M** | 0.261 | 0.018 | 0.006 | | | | | |
| **128 x 128** | **10** | **NM** | 1.431 | 0.111 | 0.045 | 0.565 | 0.153 | 0.147 | **1.148±0.147** | 8.08E-07 |
| | | **M** | 0.866 | 0.170 | 0.051 | | | | | |
| | **5** | **NM** | 1.125 | 0.040 | 0.016 | 0.453 | 0.097 | 0.083 | **0.899±0.083** | 1.65E-08 |
| | | **M** | 0.672 | 0.115 | 0.035 | | | | | |
| | **2.5** | **NM** | 0.694 | 0.043 | 0.017 | 0.246 | 0.048 | 0.051 | **0.571±0.051** | 1.84E-07 |
| | | **M** | 0.447 | 0.050 | 0.015 | | | | | |

**Supplementary Table T2.** Statistical parameters of sill values of non-metastatic (NM) and metastatic (M) tissues for different Gaussian Filter Sigma $\sigma$ ($px$). Mean sill values, Standard Deviation (StDev), Standard Error of Mean (SE Mean), the difference of M and NM mean values (D), Standard Deviation of Difference (StDev of D), 95% Confidence Interval for the difference (95% CI for D), Threshold ± 95%CI for D, P-Value statistic parameter for the hypothesis NM ≠ M.

| | Tissue | Mean | StDev | SE Mean | D | StDev of D | 95% CI for D | NM≠ M : P-Value |
|---|---|---|---|---|---|---|---|---|
| **Theta distribution skewness** | **NM** | -0.138 | 0.218 | 0.082 | 0.566 | 0.191 | 0.217 | 0.000128 |
| | **M** | 0.428 | 0.174 | 0.052 | | | | |
| **Theta distribution kurtosis** | **NM** | 1.857 | 0.178 | 0.067 | 0.339 | 0.190 | 0.194 | 0.00210 |
| | **M** | 2.195 | 0.197 | 0.059 | | | | |
| **Average z-height (*nm*)** | **NM** | 2061.7 | 486.2 | 146.6 | 102.6 | 416.1 | 376.3 | 0.569 |
| | **M** | 2164.3 | 260.2 | 98.3 | | | | |
| **RMS roughness (*nm*)** | **NM** | 689.6 | 127.4 | 38.4 | 306.0 | 130.5 | 139.6 | 0.000422 |
| | **M** | 995.6 | 135.7 | 51.3 | | | | |
| **Mean Hurst exponent of 512 lines (topo).** | **NM** | 0.860 | 0.040 | 0.013 | -0.040 | 0.037 | 0.037 | 0.0369 |
| | **M** | 0.820 | 0.031 | 0.012 | | | | |
| **Hurst exponent of 512 l lines (topo)** | **NM** | 0.603 | 0.040 | 0.013 | 0.013 | 0.039 | 0.041 | 0.509 |
| | **M** | 0.616 | 0.038 | 0.014 | | | | |
| **Phase average value (*V*)** | **NM** | 7.422 | 0.970 | 0.292 | -0.023 | 1.141 | 1.353 | 0.969 |
| | **M** | 7.399 | 1.380 | 0.522 | | | | |
| **Phase "RMS roughness" (*V*)** | **NM** | 2.703 | 0.368 | 0.111 | 0.219 | 0.381 | 0.409 | 0.265 |
| | **M** | 2.922 | 0.401 | 0.152 | | | | |
| **Mean Hurst exponent of 512 lines (Phase)** | **NM** | 0.703 | 0.084 | 0.025 | -0.060 | 0.067 | 0.058 | 0.0437 |
| | **M** | 0.643 | 0.021 | 0.008 | | | | |
| **Hurst exponent of 512 lines as one line (Phase)** | **NM** | 0.613 | 0.020 | 0.006 | 0.016 | 0.022 | 0.026 | 0.182 |
| | **M** | 0.629 | 0.026 | 0.010 | | | | |
| **F.D. Box Counting** | **NM** | 2.321 | 0.035 | 0.010 | 0.024 | 0.031 | 0.029 | 0.0976 |
| | **M** | 2.345 | 0.023 | 0.009 | | | | |
| **F.D. Partioning** | **NM** | 2.512 | 0.032 | 0.010 | -0.026 | 0.033 | 0.035 | 0.131 |
| | **M** | 2.486 | 0.034 | 0.013 | | | | |
| **F.D. Triangulation** | **NM** | 2.406 | 0.033 | 0.010 | 0.028 | 0.031 | 0.031 | 0.0762 |
| | **M** | 2.434 | 0.028 | 0.011 | | | | |
| **F.D. Power Spectrum** | **NM** | 2.168 | 0.056 | 0.017 | -0.109 | 0.054 | 0.054 | 0.000633 |
| | **M** | 2.059 | 0.049 | 0.018 | | | | |

**Supplementary Table T3**. Statistical parameters of theta distribution skewness (Fig. 6b) and kurtosis (Fig. 6c), average z-height (*nm*) (Fig. S5a), RMS roughness (*nm*) (Fig. S5b), Mean Hurst exponent of 512 lines (topography) (Fig. S5c), Hurst exponent of 512 lines as one line (topography) (Fig. S5d), phase average value (*V*) (Fig. S6a), Phase "RMS roughness" (*V*) (Fig. S6b), Mean Hurst exponent of 512 lines (phase) (Fig. S6c), Hurst exponent of 512 lines as one line (phase) (Fig. S6d), Fractal Dimension Box Counting (Fig. S7a), Partitioning (Fig. S7b), Triangulation (Fig. S7c) and Power Spectrum (Fig. S7d) methods, of non-metastatic (NM) and metastatic (M) tissues. Mean values, standard deviation (StDev), standard error of mean (SE Mean), the difference of M and NM mean values (D), standard deviation of difference (StDev of D), 95% Confidence Interval for the difference (95%CI for D), P-Value statistic parameter for the hypothesis NM ≠ M.
